# Discovery of novel G-quadruplex stabilizing compounds from medicinal plant & Evaluates their cellular toxicity

**DOI:** 10.1101/2023.07.08.548213

**Authors:** Neha, Prashant Ranjan, Parimal Das

## Abstract

G-Quadruplexes (G4Q) are higher-order, four-stranded structures that can be formed by repeated guanine tracts in human genomes. In this study, we used a structure-based virtual screening of phytomolecules derived from medicinal plants in order to discover new natural DNA G4Q binders. The top 40 ligands were sorted out based on binding affinity values after completing a docking study on 314 phytomolecule samples with parallel and mixed hybrid structure. Further Thermal melting, circular dichroism, and fluorescence displacement experiment was used as a preliminary screening tool to determine the potential stabilizing properties of β-sitosterol-β-D-glucoside, and Glabrolide. The cytotoxicity experiments were conducted on HEK293T cells and found that both of the tested phyto molecules are non-toxic for up to 150 μM concentration. Based on their cytotoxic experiments at the suggested high concentration, these phyto molecules may potentially be employed as G-Quadruplex targets in future research or applications. These results suggest that the plant may be a “lead” in the future for the development of novel therapies for diseases.

## 1. Introduction

It is now widely accepted that repeating G-tract sequences in genomic DNA can fold into higher-order G-quadruplex (G4Q) structures (Das & Verma, 2020) and that the human genome and many other genomes contain a large number of G4Q forming sequences (Huppert & Balasubramanian, 2005). They make up the tandem repeat sequences in telomeric DNA, which are found in abundance in oncogene promoter sequences. They also are present in 5’-UTR sections, introns, and various fragile/breakpoint areas in the human genome. These structures can obstruct transcription and translation if left unresolved and stabilized by small molecules, and are hence a type of gene targeting. G4Q targeting with the help of small compounds is developing as a novel strategy for cancer therapies and may have applicability in other disease types, including pathogenic and viral infections as well as repeat-expansion neurological diseases (Ohnmacht & Neidle, 2014). Recently, the direct observation of G4Q DNAs and RNAs in human cells and the discovery that these structures can be seen to be stabilized in cells by small chemicals have corroborated the general idea that G4Q DNAs and RNAs are therapeutically vulnerable hot spots (Biffi et al., 2013)(Biffi et al., 2014)(Yuan et al., 2013). The exponential growth of the G4Q field brings to light some of the key achievements and difficulties in this area, highlighting those more general concepts that are particularly relevant to the design and discovery of therapeutic G4Q-targeted small molecules. Despite the fact that there is still much to be done in the creation of effective medications that target G-quadruplexes, some encouraging lead compounds have been found. Natural alkaloids, a substantial class of natural compounds with numerous and significant biological activity, make up some of these known G4Q ligands. These substances have undergone extensive investigation for their potential as G-quadruplex ligands because they exhibit chemical diversity, structural complexity, and are accessible from natural sources (Che et al., 2018). Since the beginning of human agriculture, plants have been one of the most significant sources of medicine. Demand for plant-based drugs, food supplements, cosmetics, health products, and other products are rising. All indigenous or complementary medical systems rely heavily on medicinal plants. A new chemical molecule that can lead to a new medicine can be discovered from sources like medicinal plants (Kalia, 2009). Many phytocompounds found in plants are effective at binding to and controlling the thermal stability of the G4Q. With a view to investigating a growing number of options for a plant-based G4Q binder we discovered a new G4Q stabilizing phytomolecules derived from world wide distributed medicinal plants by means of a structure-based approach based on docking experiments against parallel and mixed hybrid G4Q structure in order to discover new natural DNA G4Q binders. 314 selected phytomolecules were subjected to docking with G-quadruplex structures, and the top 40 ligands were shortlisted based on binding affinity values. The stabilization of the G-quadruplex structure with the two phytomolecules β-sitosterol-β-D-glucoside and Glabrolide has been investigated further using *In vitro* biophysical methods. In this study, the plant *Morus alba* was found to be the source plant of the β-sitosterol-β-D-glucoside compound from our selected plants according to the database. However, alternative source plants for this phytochemical, like *Selaginella bryopteris* (Verma et al., 2015) and *Cassia fistula* (Jain et al., 2013), have been mentioned in a few earlier articles. β-sitosterol-β-D-glucoside is a phytosterol which together with β-sitosterol has so far been used for the treatment of benign prostate hypertrophy. At incredibly low concentrations, this phytosterol has also been shown to enhance the *In vitro* proliferating response of T-cells triggered by sub-optimal concentrations of phytohaemagglutinin (PHA) levels (Bouic et al., 1996). The fact that β-sitosterol-β-D-glucoside has activity against both bacterial and fungal strains while β-sitosterol only exhibits activity against a small number of bacterial stains suggests that it is a more effective antibacterial agent than β-sitosterol. This is likely due to the glucosidic moiety of the compound (Verma et al., 2015). Glabrolide is aliphatic heteropolycyclic compound that belongs to the class of organic compounds known as triterpenoids. These are terpene molecules containing six isoprene units. Glabrolide is an extremely weak basic (essentially neutral) compound based on its pKa (https://hmdb.ca/metabolites/HMDB0034515). *Glycyrrhiza glabra* one of the common traditional Chinese medicinal plants used as a flavoring and sweetening agent for tobaccos, chewing gums, candies etc. is a source plant for glabrolide on our list of chosen plants. *Glycyrrhiza glabra* is one of the most widely used herbs from the ancient medical history of Ayurveda (Kaur et al., 2013). A possible role for glabrolide against the SARS-CoV-2 proteins and as a potential disease-modifying medication for the treatment of Alzheimer’s disease (AD) by inhibiting β-Site amyloid precursor protein cleaving enzyme 1 (BACE1) have been suggested by an *In silico* drug repurposing analysis (Arif et al., 2020; Joshi et al., 2022). Studies on the biological role and involvement of both β-sitosterol-β-D-glucoside and glabrolide have been reported, but no studies have been reported for their potential role as a G-quadruplex binder. In this study, we have the first time discovered the G-quadruplex binding potential of these phytocompounds.

## 2. Materials and Methods

### 2.1 Materials and Reagents

PRCC G-quadruplex (PRCC-G4Q), sequence within the PRCC part of *PRCC-TFE3* fusion gene and telomeric G-quadruplex (Tel-G4Q) have been utilized in this study. Sequences of oligonucleotides were mentioned in appendix. All oligonucleotides were created and purchased from Eurofins Genomics India Pvt. Ltd. and utilized as received. Dulbecco’s Modified Eagle’s Medium (DMEM), fetal bovine serum (FBS), and Penicillin-Streptomycin antibiotic cocktails were purchased from Himedia. Glabrolide (Glab) were purchased from ChemFaces, China while β-sitosterol-β-D-glucoside (β-Sito) and thioflavin-T were bought from Sigma-Aldrich. All the three compounds were dissolved in DMSO for stock preparation.

### 2 Phytochemical Drug library preparation

All plant phytochemicals were retrieved from IMPPAT (Indian Medicinal Plants, Phytochemistry and Therapeutics) at “https://cb.imsc.res.in/imppat/home” and Phytochem Library at “https://phytochem.nal.usda.gov/phytochem/search/list”. PubChem (https://pubchem.ncbi.nlm.nih.gov), and Chemspider (http://www.chemspider.com), databases were used to acquire the list of chemical compounds in sdf format.

### 2.3 Retrieval of target structures of parallel and Mixed hybrid-type telomeric G-quadruplexes

Crystal structures of parallel, antiparallel and mixed hybrid-type telomeric G-quadruplexes have been retrieved from the Protein Data Bank database (Berman et al., 2007). For parallel, antiparallel and mixed hybrid, the PDB IDs were 1KF1, 143D, and 2HY9, respectively. The 3D structures thus generated were visualized by Discovery Studio 4.0 (Studio, 2008) and Chimera (Pettersen et al., 2004). Cleaning of structure such as water molecules was done same from them.

### 2.4 Molecular Docking

Molecular Docking were performed using AutoDock Vina (Trott & Olson, 2010), based on search algorithm and creates large number of potential poses of drug molecule in their coupling site. The pdb file of target DNA is converted into pdbqt format by Auto Dock Tools. Also the ligand molecule has been saved in pdbqt format via Open Bable. The grid was set of all targets. Parallel_1kf1 size X=86 size Y= 90 and size Z= 106 with centre of the grid box 23.591 × -0.549× -8.645, mixed hybrid_2hy9 size X=40 size Y= 40 and size Z= 40 with centre of the grid box 0 × 0× 0, and antiparalle_143D size X=56 size Y= 78 and size Z= 58 with centre of the grid box 0.516 × 0.422× -2.639. After approval of the docking convention, virtual screening was conducted by rigid docking into the binding site of all targets DNA. Molecular connections between protein-ligand complexes, including hydrogen bonds and non covalent and covalent interactions were analyzed and portrayed by utilizing Discovery studio and Chimera.

### 2.5 G-quadruplex Sample Preparation

In this investigation, desalted and HPSF-purified oligonucleotides were employed. In order to anneal the oligonucleotide sequences (PRCC-G4Q & Tel-G4Q) into the G-quadruplex conformation, the oligonucleotide sequences were dissolved, heated with 10 mM Tris pH 7.4 and 100 mM KCl at 95°C for 5 min in a heat block, and then gradually cooled at room temperature for overnight. They were then stored at 4°C.

### 2.6 Circular Dichroism Spectroscopy

CD spectra of folded PRCC-G4Q & Tel-G4Q (5 µM) in the presence or absence of 30 μM of β-Sito and Glab were measured in, JASCO J-1500 CD Spectrophotometer fitted with a thermostable cell holder using a quartz cuvette with a 2.0 mm path length. Spectra were recorded at 220-320 nm with 50 nm min-1 scanning speed and 1.0 nm bandwidth. The data interval time was set to 1 sec and the data pitch was 0.2 nm. Each spectrum was captured three times, smoothed, and the baseline removed. The study’s buffer consisted of 10 mM Tris and 100 mM KCl at pH 7.4.

### 2.7 Thermal Melting Study

Using the CD melting method, the thermal stability of both PRCC & Telomeric G-quadruplexes DNA in the absence or presence of both compounds Glab and β-Sito was investigated. 5 µM of both PRCC-G4Q & Tel-G4Q were annealed by heating with 10 mM Tris pH 7.4 and 100 mM KCl at 95°C for 5 min followed by slow cooling to room temperature. Thermal melting of G-quadruplex samples was observed at 295 nm by heating the samples from 30°C to 90°C at a heating rate of 1°C min^-1^. The formula applied to calculate the fraction folded was Qt-Qu/Qf-Qu, where Qt refers to the molar ellipticity at temperature T and Qu and Qf are the molar ellipticities of the fully unfolded (highest temperature) and fully folded (lowest temperature), respectively. Thermal melting study was carried out by JASCO J-1500 CD spectrophotometer equipped with a Peltier temperature controller.

### 2.8 Fluorescence Intercalating Displacement Assay

Both PRCC and Telomeric oligonucleotides, bearing G-quadruplex sequence tend to fold into G-quadruplex structure as mentioned in the above section. 5 µM of G-quadruplex oligos were combined with 5 µM of thiazole orange (TO) in a 1:1 ratio and equilibrated for 2-3 hours. Fluorescence spectra of solution of TO with G-quadruplex oligos in the absence or presence of Glab and β-Sito with increasing concentration (0.5-40 μM) recorded at 25°C using A Perkin Elmer LS55 fluorescence spectrometer. Excitation and emission slits for fluorescence measurements were 10 and 5 nm respectively and the wavelengths for the excitation and emission of TO were set to 501 nm and 500-700 nm, respectively.

### 2.9 MTT assay

MTT assay was performed according to Neha *et al*., 2023 (Neha et al., 2023). In brief HEK293T cells were cultivated in DMEM culture media within the humidified incubator maintained with 5% (v/v) CO2. 10% FBS and an antibiotic cocktail containing 100 g/ml streptomycin and 100 U/ml penicillin were added to the culture media. For 24 hours, cells were grown in 96-well plates at a density of 2×10^4^. The next day, cells were subjected to 24 hours of exposure to both phytomolecules at different concentrations (25-150 μM). After 24 hours, the media was changed to DMEM with 0.5 mg/ml MTT and incubated at 37°C for 4 hours. In order to dissolve the blue/purple formazan, the MTT solution was removed after 4 hours, 100 μl of DMSO was added, and the mixture was then incubated in the dark for 10 minutes. The purple color end product formazan was quantified using a microplate reader (Bio-Rad) to measure absorbance at 570 nm, and the IC50 dose was calculated using a dose-response linear regression curve.

## 3. Results and Discussion

### 3.1 Phytochemical library

We identified 1226 chemical compounds in total from the database, and further polysaccharides, fatty acids, and essential oils were excluded. Additionally, we ignored molecules that weren’t found in databases and took duplicate compounds out of the library. We finally acquired the phyto drug library, which contains 314 phyto-compounds. List of selected medicinal plants and shortlisted phytochemical for the docking analysis were enlisted in supplementary file (S1).

### 3.2 Docking study

Binding energy of all docked phytomolecules were calculated and top 40 phyochemicals on the basis of binding affinity for both parallel anti antiparallel G-quadruplex were screened out supplementary file (S1). Among this top 40 phytochemical list two phytomolecules; β-sitosterol-β-D-glucoside (β-Sito) and glabrolide (Glab) were selected for further *In vitro* analysis. Binding affinity of docked phytomolecules were enlisted in supplementary file (S1). The binding scores of the parallel G-quadruplex with β-Sito & Glab was -16.8 and -10.8 respectively and of antiparallel G-quadruplex with β-Sito & Glab was -14.6 and -11.5 respectively. However, β-Sito & Glab revealed binding energy of -6.6 and -8.1 respectively with mixed hybrid G-quadruplex. The docked complex’s noncovalent and covalent interactions are shown in Figure 1. β-Sito interaction with parallel quadruplexes revealed hydrogen bond, Pi (π)-sigma, π-Alkyl, and van der waal force attraction. While Interaction with Glab revealed π-sigma interaction. β-Sito interaction with mixed hybrid quadruplex shows π-Alkyl, covalent, and vander waals bonding. However, with Glab revealed van der waal force of attraction and carbon hydrogen. Interaction pattern of parallel quadruplexes with β-Sito and Glab suggest end stacking interaction while mixed hybrid showed intercalating and groove binding patterns (Figure 2).

**Figure 1:**
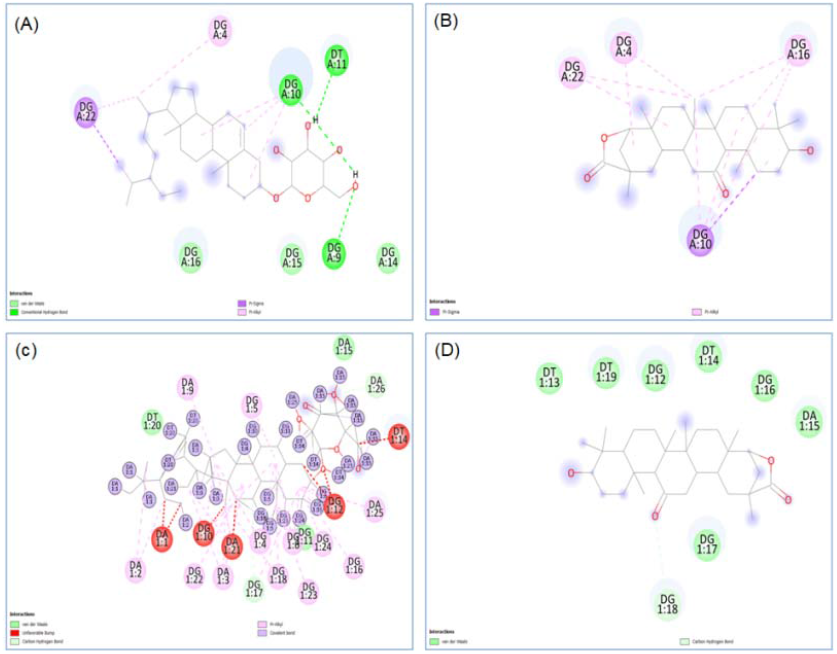
Annotation of 2D interactions of Phyto-molecules with parallel and mixed hybrid G-quadruplex obtained with Autodock Vina. A. Parallel G-quadruplex_β-Sito, B. Parallel G-quadruplex_Glab, C. Mixed hybrid G-quadruplex_β-Sito, D. Mixed hybrid G-quadruplex_Glab. Different colour indicates the way of interaction between the DNA and molecules

**Figure 2:**
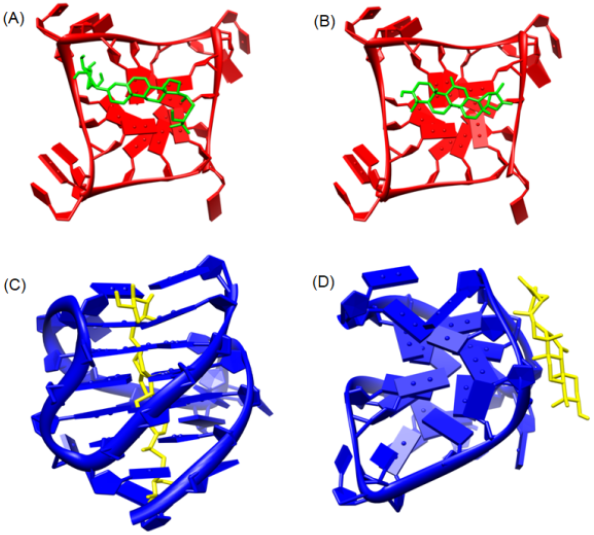
Schematic 3D representation of the interaction pattern of the DNA-Ligand complex. Parallel G-quadruplex interaction with β-Sito & Glab (A & B), Mixed hybrid type G-quadruplexes interaction with β-Sito & Glab (C & D). The end-stacked interaction pattern was observed with parallel quadruplexes. However, intercalating and groove binding patterns were seen with hybrid-type telomeric G-quadruplexes. Yellow colour and green color indicate ligand molecules.

### 3.3 G-quadruplex formation

The first step was to get CD spectra to verify that both oligo samples were folded into G-quadruplex conformation. A hybrid-type mixed topology was visible in the CD spectra of Tel-G4Q (human telomere G-quadruplex DNA), which had shoulder peaks at 265 nm with negative and positive peaks at 240 and 290 nm respectively. The CD spectra of PRCC-G4Q, on the other hand, contain two peaks that are suggestive of parallel-stranded G4Q topologies: a negative peak at 240 nm and a positive peak near 263 nm (Figure 3). However, peak intensities change in both G4Qs, indicating that each phytomolecules interacts with the G4Q structure, despite the fact that the parallel and mixed hybrid topologies of PRCC-G4Q and Tel-G4Q, respectively, were still maintained after incubation with both phytomolecules independently (Figure 4).

**Figure 3:**
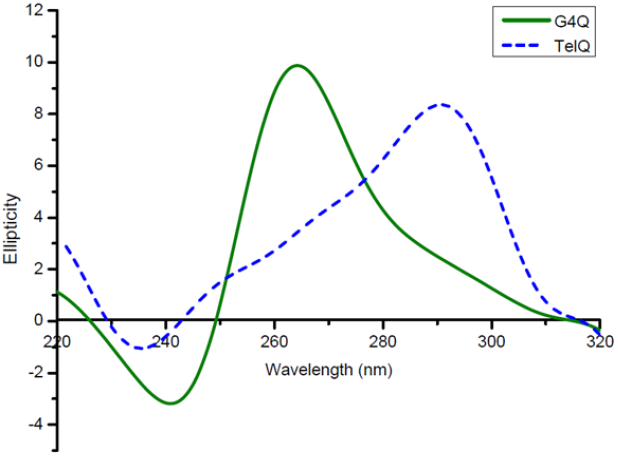
CD spectra of the human telomeric (Tel-G4Q) sequence and the PRCC-G4Q (G-rich sequence within the PRCC stretch of the *PRCC-TFE3* fusion gene). Illustrating parallel G-quadruplex structure creation in PRCC-G4Q and mixed hybrid G-quadruplex formation in Tel-G4Q

**Figure 4:**
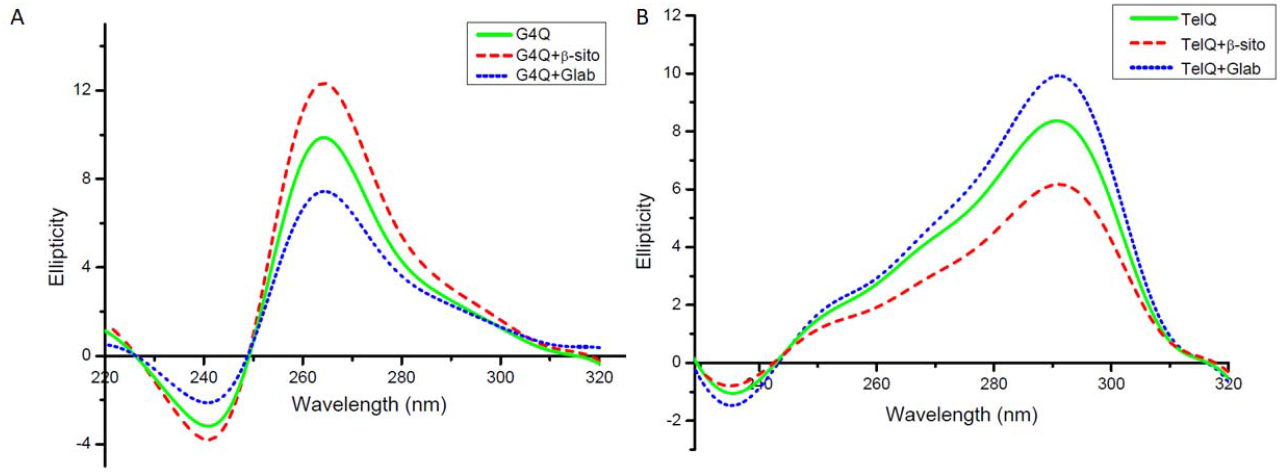
In the presence of 30 μM β-Sito and Glab, the CD spectra of PRCC-G4Q and Tel-G4Q. Illustrating how the presence of both β-Sito and Glab causes the formation and maintenance of parallel G-quadruplex structure in PRCC-G4Q (I) and mixed hybrid G-quadruplex structure in Tel-G4Q (II)

### 3.4 G-quadruplex thermal stabilization

The thermal stabilization of both of the selected G-quadruplex oligos in the presence of glabrolide and β-sitosterol was examined using the CD melting method. The determination and comparison of each melting temperature (Tm) transitions were made accessible by plotting the fraction-folded vs. temperature graph. When combined with β-Sito and Glab, the Tm of PRCC-G4Q increased to 65.5°C and 63.9°C, respectively. Similar results were seen with Tm of Tel-G4Q, which increased from 58.7°C to 64°C with β-Sito and 62°C with Glab (Figure 5).

**Figure 5:**
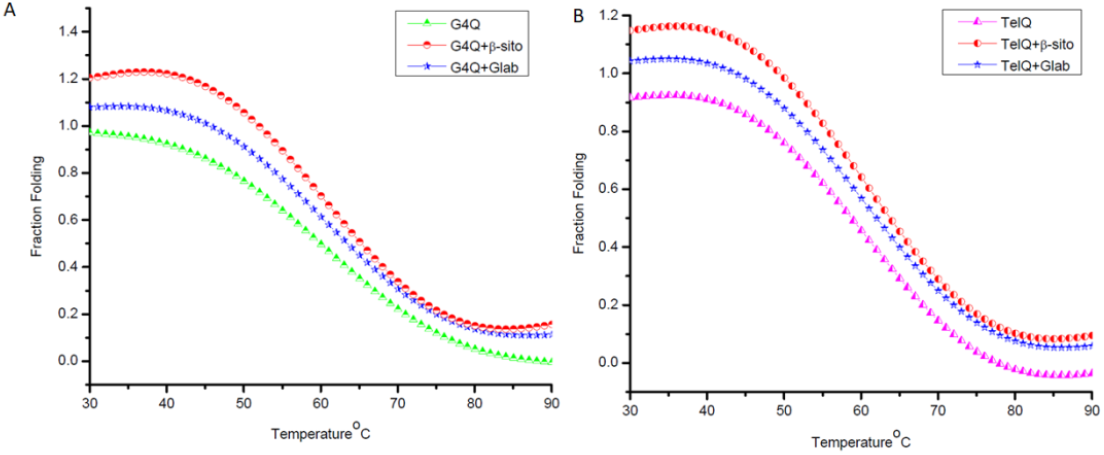
CD melting curve of 5 μM PRCC-G4Q and Tel-G4Q in the presence of 10 μM of β-Sito and Glab. (I) Showing increment in melting temperature (Tm) of PRCC-G4Q in the presence of both β-Sito and Glab. (II) Showing increment in melting temperature (Tm) of Tel-G4Q in the presence of both β-Sito and Glab

### 3.5 Fluorescence Intercalating Displacement assay

Graphing the proportion of TO displacement vs. the concentration of specific phytomolecule resulted in the formation of FID curves (Figure 6). The percentage displacement (PD) was calculated from the fluorescence area (FA) using the equation PD = 100 -[(FA/FA_0_) 100]. The areas of fluorescence with and without phytomolecule added are designated FA and FA_0_, respectively. The concentrations of phytomolecules required for the 50% displacement of TO occur (DC50), were identified from the FID plot using a linear regression fitting curve. In both phytomolecules, the DC50 value for the two G4Qs samples was found to be marginally higher. While the DC50 concentration of Glab was 130 and 200 μM for PRCC-G4Q and Tel-G4Q, respectively, the DC50 concentration of β-Sito was discovered to be around 113 and 114 μM for PRCC-G4Q and Tel-G4Q, respectively.

**Figure 6:**
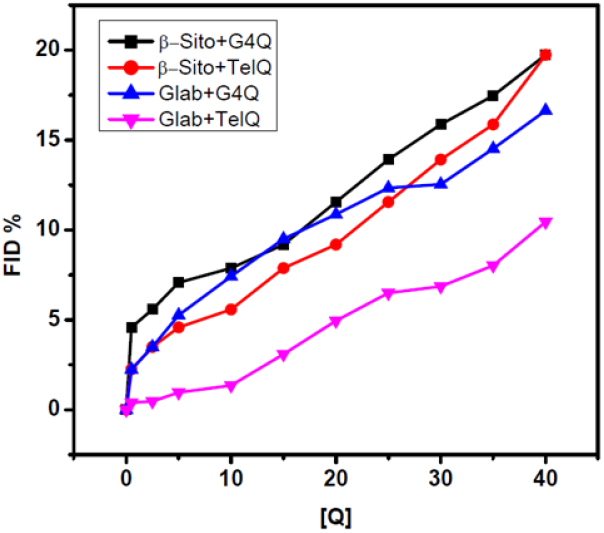
FID curve of both β-Sito and Glab with increasing concentration of PRCC-G4Q (I), and Tel-G4Q (II) obtained by plotting %FID vs. concentration

### 3.6 β-Sito and Glab are nontoxic towards HEK293T cells

It is interesting to investigate the toxicity of these phytomolecules towards HEK293T cells given their potential value as G-quadruplex targeting ligands in a cellular environment. In this case, the toxicity of these phytomolecules on HEK293T cells was assessed using the MTT assay. Both phytomolecules were applied to cells at progressively higher concentrations, up to a maximum of 150 μM. Although the maximum cell death percentage at a concentration of 150 μM was only found to be around 30%, Figure 7 shows a graph of cell death percentage vs. phytomolecule concentration. The IC50 values for β-Sito and Glab were found to be 247 and 215 μM, respectively.

**Figure 7:**
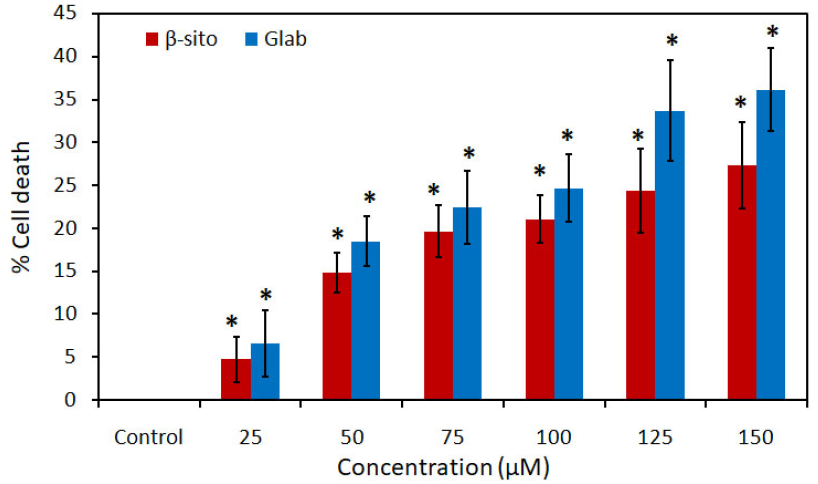
MTT graph showing antiproliferative effect of both phytomolecules β-Sito and Glab on HEK293T cell line. The error bar shows the mean and SD. The paired t-test was used to measure statistical significance. *P ≤ 0.05 compared with control vs. treatment groups

4. Discussion

Due to the fact that allopathic medications have greater side effects, there has been a surge in demand for phytopharmaceuticals worldwide. This provides a solid foundation for choosing which plant to study further in terms of phytochemistry and pharmacology. Chemical components suggest that the plant may be used as a “lead” in the future for the development of novel therapies for diseases. Further research needs to be conducted in this area to explore the increasing availability of phytomolecules for potential therapeutic uses. We discovered a new G4Q stabilizing molecule by conducting a structure-based virtual screening of phytomolecules derived from medicinal plants in order to discover new natural DNA G4Q binders. In this study 41 medicinal plants with a global distribution have been chosen. Since many years ago, these plant components have been utilized to make traditional Ayurvedic, herbal, and natural remedies (Andrighetti-Fröhner et al., 2005; Bagla et al., 2012; McCutcheon et al., 1995; Mukhtar et al., 2008; P. Ranjan et al., 2020; Semple et al., 1998). The top 40 molecules were shortlisted based on binding affinity after completing a docking study on 314 chosen phytomolecules with two different G-quadruplex topologies; parallel and anti parallel. We have chosen two phytomolecules (β-sitosterol-β-D-glucoside and glabrolide) from this list of the top 40 phytocompounds for additional *In vitro* examination. We were interested in seeing the *In vitro* binding ability of molecules having lower binding affinity in virtual screening in addition to molecules showing higher binding affinity, which is why we have selected these two molecules since one is ranked one in the top 40 list and the other is ranked 40. When measured for binding affinity, β-Sito placed first with antiparallel G4Q with a value of -14.6 and second with parallel G4Q with a value of -16.8. Glab, on the other hand, ranked 10^th^ in antiparallel G4Q with an affinity value of -11.5 and 40^th^ in parallel G4Q with a binding affinity of -10.8. Interestingly, when β-Sito and Glab were docked with mixed hybrid G-quadruplex, the resulting binding energies were lower than parallel and antiparallel which suggests comparatively weaker binding affinities with mixed hybrid G-quadruplex conformation. However it is important to consider that binding energy alone cannot explain the complete binding behavior observed *In vitro*. The binding behavior of ligands with different types of G-quadruplexes can be influenced by various factors, including the structural characteristics of the G-quadruplex and the specific interactions involved in the binding process. While binding energy is an important factor, it is not the sole determinant of the binding affinity or stability of a complex (Bulusu & Desiraju, 2020; Jena et al., 2022; Sauer et al., 1994; Sbongile & ES Soliman, 2015; Wagner & Schreiner, 2015). Mixed hybrid G-quadruplex conformation is a more complex structure than the parallel and antiparallel G-quadruplexes. The presence of mixed quadruplexes introduces additional structural heterogeneity and potentially alters the binding sites available for the ligands. This structural complexity may contribute to the observed weaker binding energies for β-Sito and Glab with the mixed hybrid topology. Interaction between ligands and receptor creates the prospect of inhibition of ligands to receptor protein. Results of molecular docking predict the possibilities of any biochemical reactions. Molecular Docking is performed to get a result on the basis of probable optimal orientations and conformations of drug molecule at groove of binding site. β-Sito found to interact with parallel G-quadruplex structure via several non-covalent interactions. These include hydrogen bonding, π-sigma interactions, π-Alkyl interactions, and van der Waals forces of attraction. Hydrogen bonding plays a crucial role in stabilizing the ligand-receptor complex through the formation of strong electrostatic interactions (Bulusu & Desiraju, 2020). π-sigma and π-Alkyl interactions involve the stacking of aromatic rings of β-Sito with the parallel quadruplex, contributing to the overall stability of the complex. Additionally, van der Waals forces, such as London dispersion forces, are responsible for attractive interactions between non-polar atoms and groups (Wagner & Schreiner, 2015). In the case of Glab, the interaction with parallel quadruplexes primarily involves π-sigma interactions. These interactions occur between the aromatic rings of Glab and the quadruplex, contributing to the binding affinity. π-sigma interactions are known for their involvement in the stacking between aromatic systems and polarizable electron-rich regions of a molecule (Jena et al., 2022). When β-Sito interacts with mixed quadruplexes, a combination of noncovalent and covalent interactions is observed which suggests a more intricate binding mechanism. Covalent interactions, such as the formation of chemical bonds, can enhance the stability of the complex. On the other hand, non-covalent interactions, like hydrogen bonding or van der Waals forces, can contribute to the overall binding affinity. Covalent interactions can occur through the formation of covalent bonds, such as carbon-hydrogen (C-H) bonds, leading to a more stable complex (Sbongile & ES Soliman, 2015). In the case of Glab interacting with mixed hybrid quadruplex, the primary non-covalent interaction observed is van der Waals forces of attraction. Van der Waals forces, specifically London dispersion forces, contribute to the attractive interactions between glabrolide and the quadruplex. These interactions play a role in stabilizing the complex (Sauer et al., 1994). Therefore, while the binding energy may provide some insights into the relative strength of the interaction, it is important to consider the overall picture, including the presence of covalent and non-covalent interactions, to better understand the binding behavior of compounds with target structure. When ligands interact with parallel quadruplexes, an end-stacked interaction pattern is observed. In this pattern, the ligand molecule is positioned at the end of the quadruplex structure, forming strong interactions with the terminal G-tetrads. This end-stacked arrangement facilitates the stacking of the ligand’s aromatic rings with the G-tetrads, resulting in a stable complex (Patel et al., 2007). In contrast to parallel quadruplexes, mixed hybrid-type G-quadruplexes exhibit two primary interacting patterns: intercalating and groove binding. In the intercalating pattern, the ligand molecule inserts itself between the G-tetrads of the quadruplex structure, forming π-π stacking interactions. This arrangement allows the ligand to intercalate into the DNA structure, resulting in a stable complex (Vianney & Weisz, 2022). The groove binding pattern involves the ligand molecule binding to the grooves formed between the G-tetrads in the quadruplex structure. This binding mode utilizes a combination of noncovalent interactions, such as hydrogen bonding, electrostatic interactions, and van der Waals forces, to stabilize the complex. The interplay of various factors, including binding energy, covalent and non-covalent interactions, and structural characteristics, collectively determines the observed binding behavior of β-Sito and Glab with different G-quadruplex structures. Further *In vitro* biophysical characterization was done to determine the potential stabilizing properties of phytomolecules β-Sito and Glab. Circular dichroism (CD) melting experiment was used as a preliminary screening tool. Two G4Q-forming oligonucleotides were employed in this work. One is the 27 bp-long PRCC G4Q sequence found in exon 1 of *PRCC* gene and the 5’ end of the *PRCC-TFE3* fusion gene (Das & Verma, 2020; Neha & Das, 2023), and the other is a 25 bp-long sequence found in human telomeric sequences. CD spectra were initially acquired to determine the conformation of each G-quadruplex architecture in both oligo samples. According to CD spectra of folded oligo samples mixed hybrid topology is formed in human telomeric sequences, while parallel G-quadruplex conformation is created in PRCC G4Q sequence. Both of these G4Q were still maintained in their respective oligo sequences following their interaction with both phytomolecules. Peak intensities do alter in the presence of each specific phytomolecule, despite the fact that no meaningful variations in the CD profiles were found for any of the analyzed G4Qs, demonstrating that each G4Q architecture is generally preserved following phytomolecule incubation. Then, by measuring the compound-induced change in the melting temperature (Tm) of G4Qs, CD-melting experiments were used to assess the stabilizing potential of β-Sito and Glab for the G-quadruplex structure. The temperature at which a complex melts and at which point it is 50% dissociated, is the midway of a certain transition is called melting temperature. The Tm value essentially provides an overall assessment of the dominant form. When temperatures are below the Tm, the structure is primarily folded, and when temperatures are above the Tm, it is mostly unfolded (Mergny & Lacroix, 2009). When β-Sito was added to the G4Qs samples, the CD melting experiment revealed an increase in Tm of around 5-6°C, compared to a slight increase in Tm (about 3°C) when Glab was applied. This clearly shows that β-Sito has a better G4Q-stabilizing impact than Glab. The behavior of a ligand in binding to DNA is frequently investigated using FID. The FID test is a simple and effective method for identifying the binding affinity of various ligands for quadruplexes. This test relies on the displacement of the fluorescent dye thiazole orange (TO) from DNA after the addition of a candidate molecule at increasing concentrations. The quantification of the binding constant for substances is made possible by the competitive displacement of the bound probe by ligands (Monchaud et al., 2008). On the two outer quartets of a G-quadruplex, the dye thiazole orange end-stacks with a strong affinity. The displacement of a specific ligand can be easily tracked by a decrease in TO’s fluorescence because TO is completely non fluorescent when free in solution but strongly fluorescent when coupled to G-quadruplex DNA (Tran et al., 2011). To learn more about the affinity of phytomolecules for G4Qs, FID tests were carried out. By calculating the DC50 value, which establishes the ligand concentration necessary to remove 50% of the TO from G-quadruplex DNA, the ligand-induces TO displacement was quantified. This value was then used to sort the ligands. The DC50 value of these two phytomolecules is quite high, which is much higher than the earlier reported G-quadruplex ligands (Pagano et al., 2018; N. Ranjan & Arya, 2013), which raises the question of whether it is safe to use these compounds in such large amounts for *In vitro* or cellular application. In order to do this, we tested the cytotoxicity of these phytomolecules on HEK293T cells and discovered that both of the tested phytomolecules are non-toxic for up to 150 μM concentration. Additionally, the IC50 concentrations for Glab and β-Sito are found to be around 215 and 247 μM, respectively, which is comparable to the previously reported IC50 concentration of β-Sito on human cell lines (Hasibuan et al., 2017; Vo et al., 2020). These phytomolecules could potentially be employed as G-quadruplex ligands in future G-quadruplex-related research or applications based on these cytotoxicity studies at the indicated high concentration. Given that these compounds are derived from medicinal plants, there may be a chance to reduce the likelihood of their side effects in comparison. Additionally it is discovered that the β-Sito phytomolecule has a greater potential binding probability with the G4Qs structure than the Glab, suggesting that phytosterol to be better G-quadruplex binder compared to aliphatic heterocyclic compounds.

## 5. Conclusion

The research and development of new drugs can benefit greatly from heterocyclic phytomolecules and natural alkaloids. Discovering putative quadruplex ligands with strong selectivity for G-quadruplexes is made possible by the polymorphism and structural complexity of phytochemicals. In order to find new natural DNA G-quadruplex binders, a structure-based virtual screening of phytomolecules produced from medicinal plants was used in this study to identify a new G-quadruplex stabilizing molecule. Further *In vitro* binding studies were conducted on two phytochemicals, including phytosterol which is top-ranked in best hit, and the 40th-ranked aliphatic heteropolycyclic molecule. Based on their cytotoxicity experiments at the suggested high concentration, these phytomolecules may be used as G-quadruplex ligands in upcoming G-quadruplex studies or applications.

## 6. Acknowledgement

We acknowledge Central Discovery Centre (CDC), Banaras Hindu University, Varanasi for Circular Dichroism facility. We acknowledge Dr. D.S Pandey, Professor, Department of Chemistry, Banaras Hindu University for providing Spectroscopy facility. We acknowledge Dr. Gopeshwar Narayan, Professor, Molecular and Human Genetics, Banaras Hindu University for providing facility of microplate reader.

## Appendix

i. PRCC-G4Q: 5’-GTTGGGGAGGGACTGGGATTGGGGTTG-3’
ii. Tel-G4Q: 5’-AGGGTTAGGGTTAGGGTTAGGG-3’

### Funding

This work was supported by a RET fellowship from Banaras Hindu University and the Institute of Eminence, BHU

### Compliance with ethical standards

### Competing financial interests

The authors declare no competing financial interests.

### Conflict of interest

The authors declare that they have no conflict of interest.

## Notes

### Competing Interest Statement

The authors have declared no competing interest.

