## Supplemental file S1 for "Discovery of novel G-quadruplex stabilizing compounds from medicinal plant & Evaluates their cellular toxicity"

**Supplementary File (S1):** Depictions of non covalent interactions pattern in details of all molecules with parallel quadruplexes and hybrid-type telomeric G-quadruplexes.

I. **Parallel G-quadruplex with ligand A**

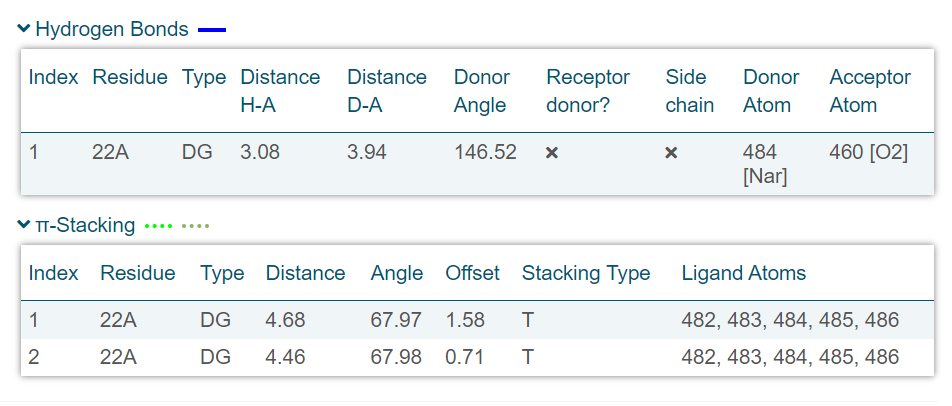

II. **Parallel G-quadruplex with ligand C**

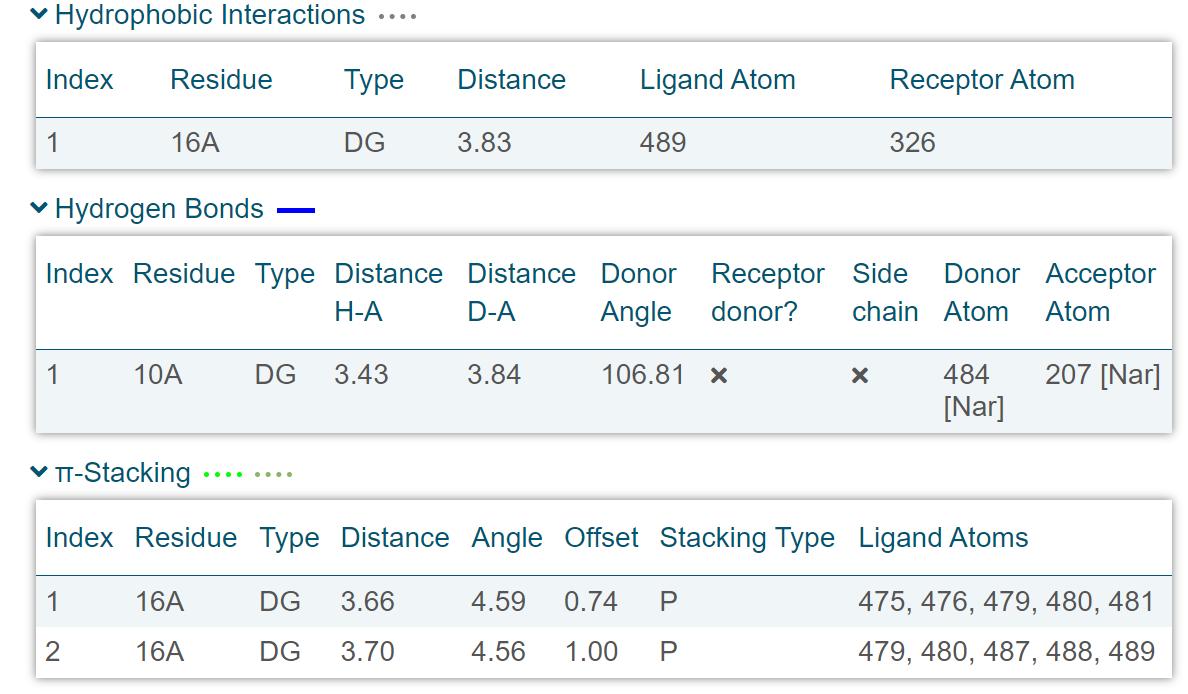

III. **Parallel G-quadruplex with complex B**

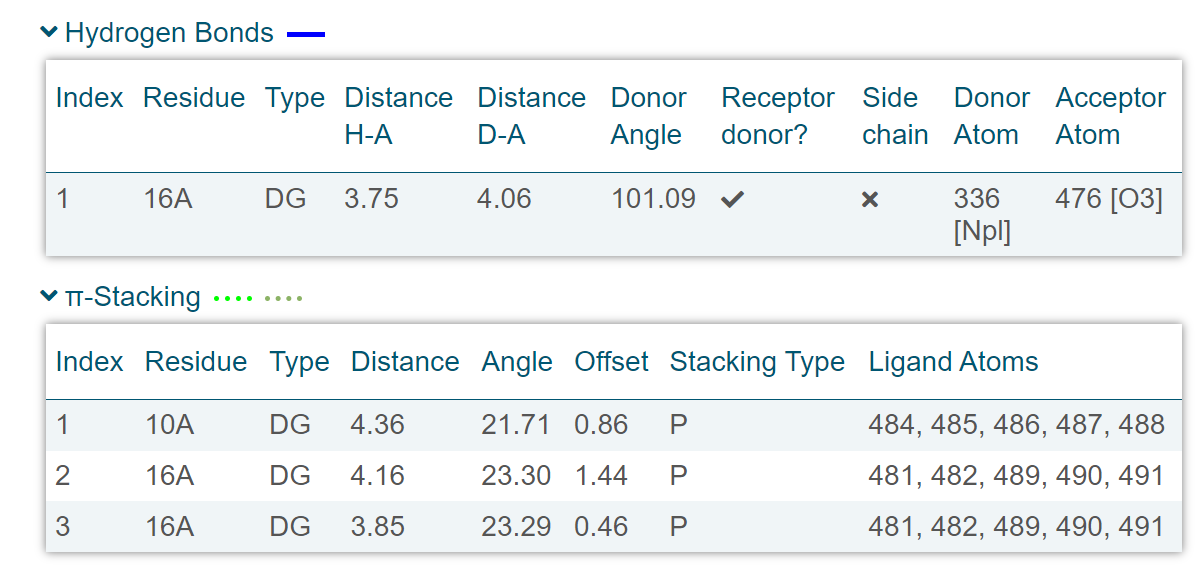

IV. **Parallel G-quadruplex with complex D**

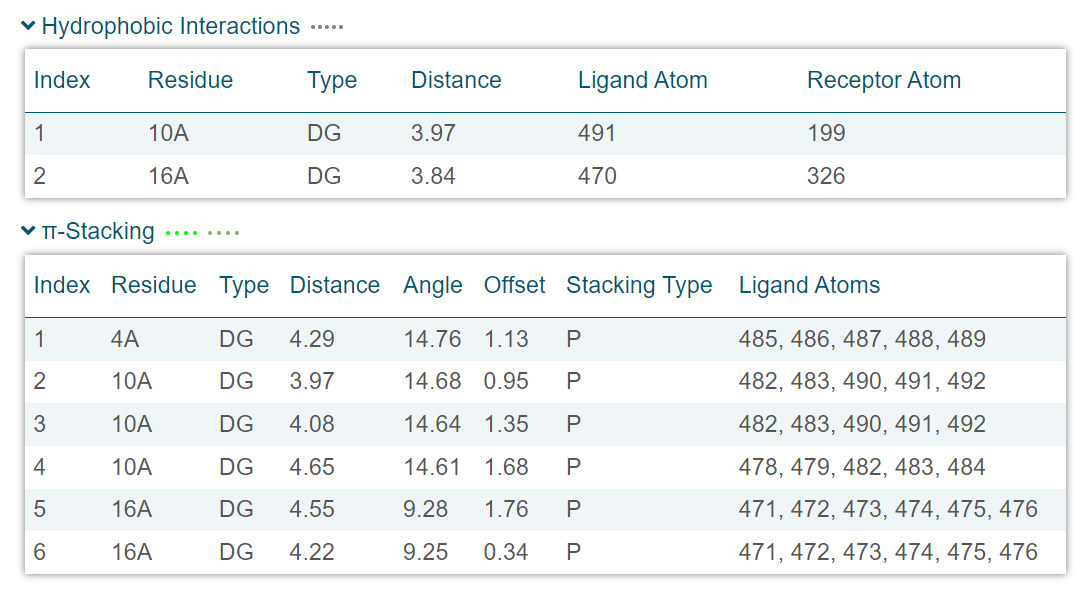

V. **Mixed hybrid type G-quadruplex with ligand A**

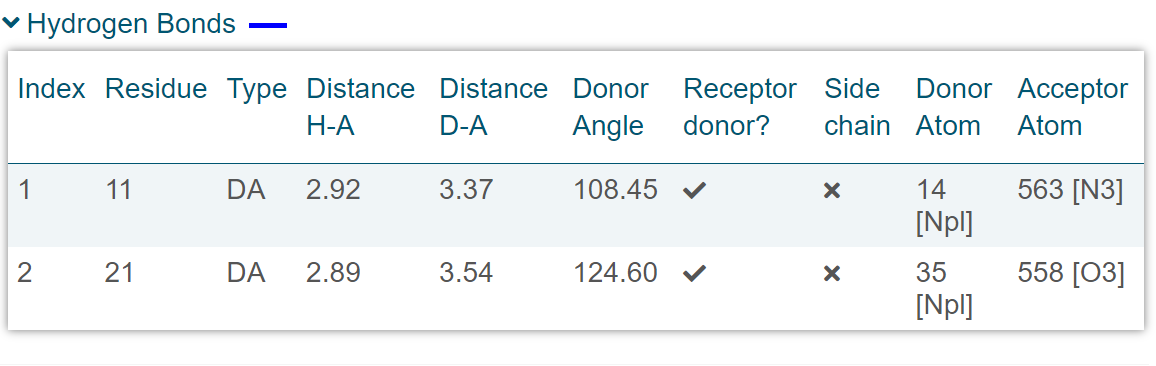

VI. **Mixed hybrid type G-quadruplex with ligand C**

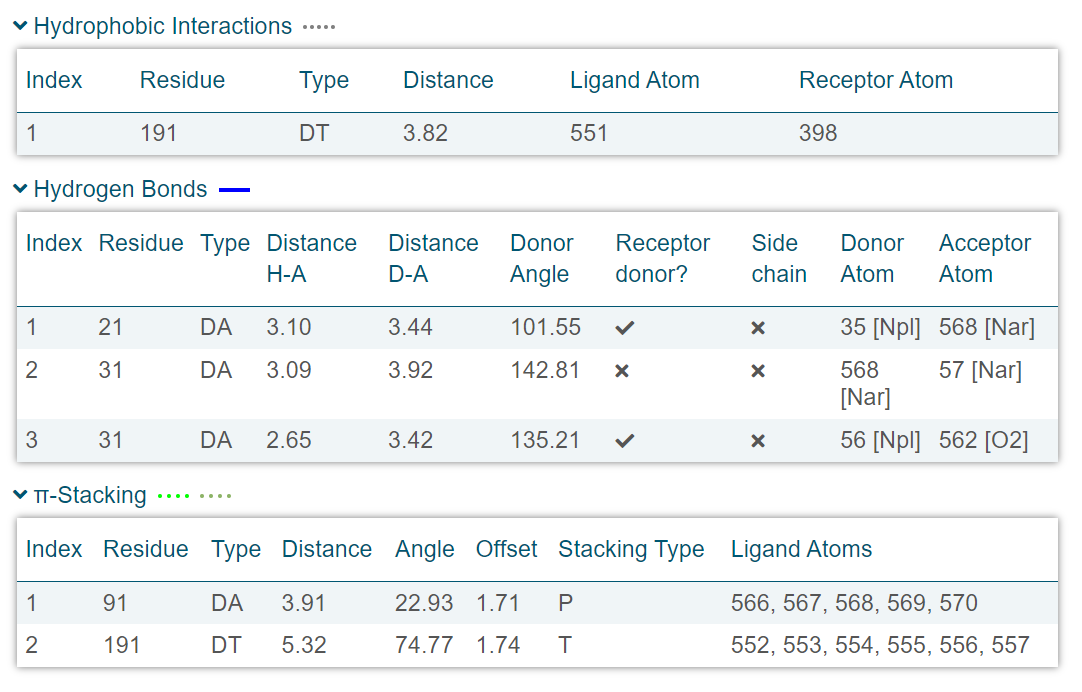

VII. **Mixed hybrid type G-quadruplex with complex B**

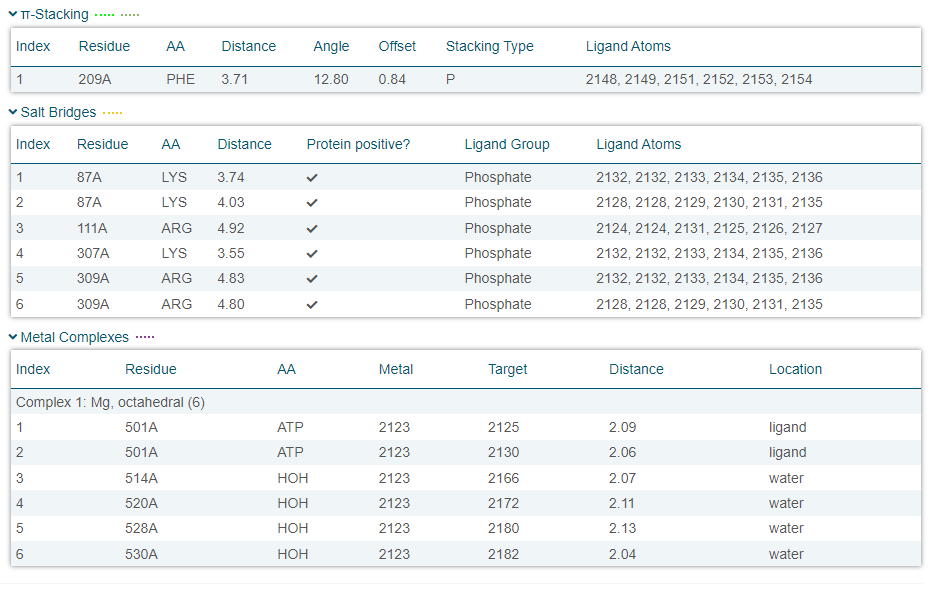

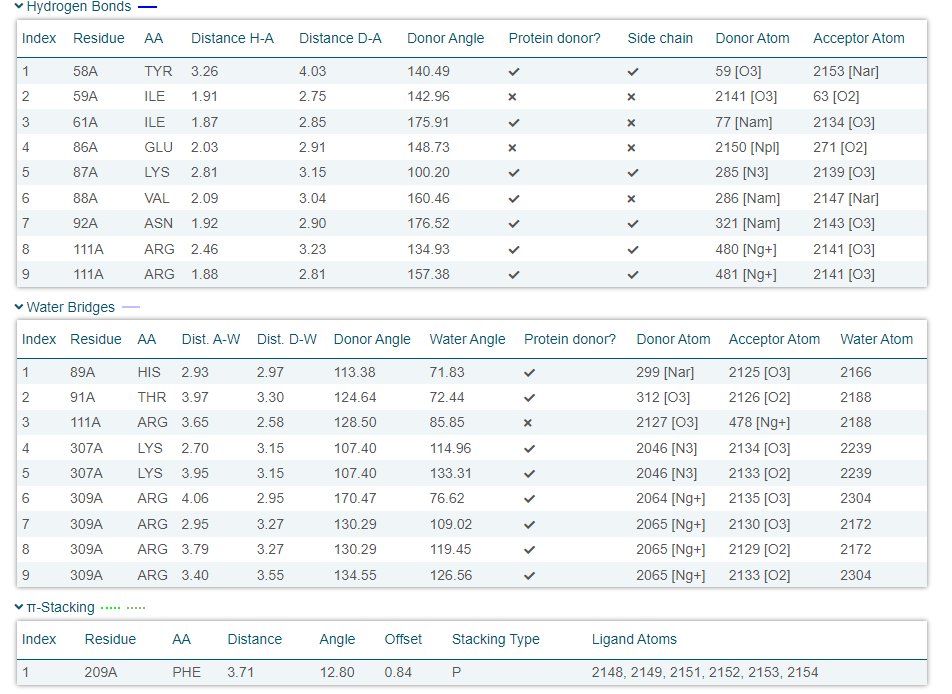

VIII. **Mixed hybrid type G-quadruplex with complex D**

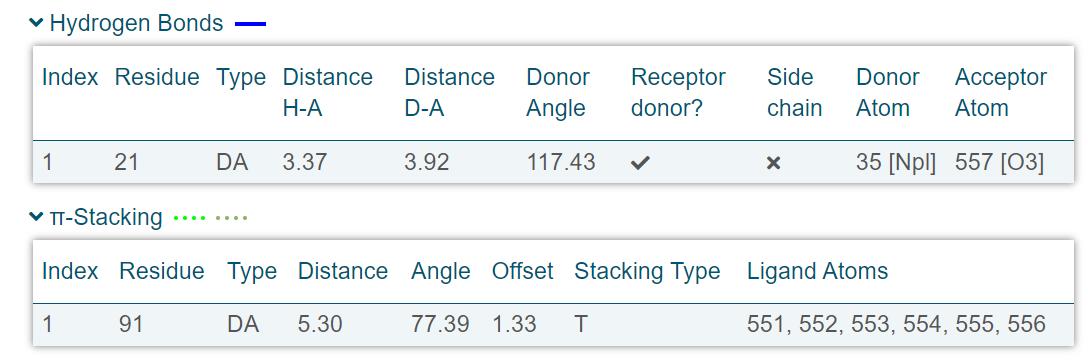
